# Rapid molecular phenotypic antimicrobial susceptibility test for *Neisseria gonorrhoeae* based on propidium monoazide viability PCR

**DOI:** 10.1101/2023.03.01.530513

**Authors:** Kristel C. Tjandra, Nikhil Ram-Mohan, Ryuichiro Abe, Tza-Huei Wang, Samuel Yang

**Affiliations:** Department of Emergency Medicine, Stanford University School of Medicine, Palo Alto, CA, USA; Departments of Mechanical Engineering and Biomedical Engineering, The Johns Hopkins University, Baltimore, Maryland, USA

**Keywords:** Neisseria gonorrhoeae, sexually transmitted disease, diagnostic methods, nucleic acid amplification technique, antimicrobial resistance

## Abstract

*Neisseria gonorrhoeae* (NG) is an urgent threat to antimicrobial resistance (AMR) worldwide. NG has acquired rapid resistance to all previously recommended treatments leaving ceftriaxone monotherapy as the first and last line of therapy for uncomplicated NG. The ability to rapidly determine susceptibility, which is currently nonexistent for NG, has been proposed as a strategy to preserve ceftriaxone by using alternative treatments. Herein, we used a DNA-intercalating dye in combination with NG-specific primers/probes to generate qPCR cycle threshold (Ct) values at different concentrations of 2 NG-relevant antimicrobials. Our proof of concept dual-antimicrobial logistic regression model based on the differential Ct measurements achieved an AUC of 0.93 with a categorical agreement for susceptibility of 84.6%. When surveying the performance against each antimicrobial separately, the model predicted 90% and 75% susceptible and resistant strains respectively to ceftriaxone and 66.7% and 83.3% susceptible and resistant strains respectively to ciprofloxacin. We further validated the model against the individual replicates and determined the accuracy of the model in classifying susceptibility agnostic of the inoculum size. We demonstrated a novel PCR-based approach to determine phenotypic ciprofloxacin and ceftriaxone susceptibility information for NG with reasonable accuracy in under 30 min, a significant improvement compared to the conventional method which takes 3 days.

**Table of Content Graphic:** 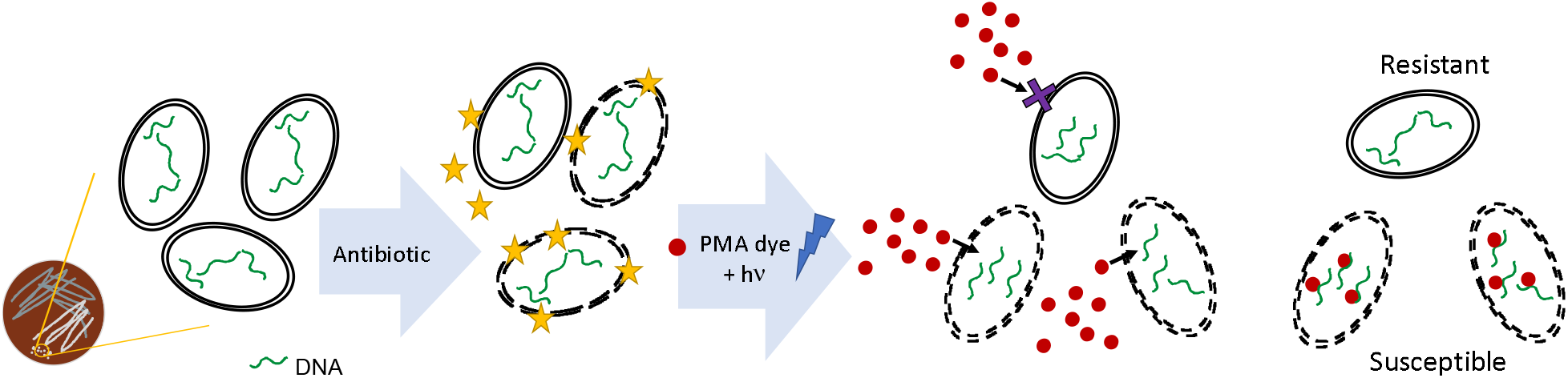

## Introduction

Antimicrobial resistance (AMR) spread in *Neisseria gonorrhoeae* (NG) is a top-tier global health threat ^1^. The World Health Organization estimates that around 87 million sexually transmitted infections each year are caused by NG ^2^. Complications from NG may include pelvic inflammatory disease, ectopic pregnancy, and infertility. Furthermore, they can facilitate the transmission of the human immunodeficiency virus (HIV) ^3^. The economic burden due to NG is substantial. In 10 years of assessment, a total of 1.2 million additional infections are suspected to occur in the US alone. This number of cases equals USD 378.2 million in predicted healthcare costs ^4^.

The CDC publishes treatment guidelines on antimicrobial selection and dosing based on susceptibility data generated by the national CDC Gonococcal Isolate Surveillance Project (GISP) ^5^. However, these susceptibility data rely on a culture-based method that is complicated to set up, labor-intensive, and takes 24-48 hours to complete ^6^. The availability of a rapid, sensitive, and automated nucleic acid amplification test (NAAT) for routine NG detection has increasingly disincentivized the use of culture-based analyses in most clinical laboratories, leading to the lack of phenotypic AMR data ^6^. In the absence of these data, patients continue to receive empirical treatment which leads to treatment failure, complications, transmission, and development of AMR.

Due to its natural competency to adapt through mutation and horizontal gene transfers ^7^, NG has acquired rapid resistance to all previously recommended treatments ^8,9^. Ciprofloxacin, which was the first-line treatment for NG since 1996, was discontinued in 2006 due to a rising rate of resistance. In 2012, dual treatment with azithromycin and ceftriaxone was introduced. Yet, a significant increase in azithromycin resistance worldwide and the identification of a very high-level resistance (> 512 μg/mL) has prompted the reversal of this recommendation ^9–11^. At the end of 2020, ceftriaxone monotherapy was decided as the recommended therapy for non-complicated gonorrhoea in the US ^12^.

Compounded by a reduced “pipeline” for new antimicrobial development, the effort to prevent NG spread is desperate. Both the CDC and WHO have labelled NG as a top-tier AMR threat. Increase in ceftriaxone resistance has been reported in Asia, Australia, and the UK ^13,14^. With only ceftriaxone as the first and last line NG therapy, rapid and reliable susceptibility test is needed to monitor emergence of NG strains with decreased susceptibility to ceftriaxone and anticipate potential outbreaks.

A rapid, point of care antimicrobial susceptibility test (AST) for NG is crucial to yield the greatest potential impact on more precise treatment selection while extending the lifeline of existing treatments ^15^. Despite the need, there is currently no clinically available NG AST that could concurrently provide rapid and reliable antimicrobial susceptibility information for both ceftriaxone and ciprofloxacin in a single platform to guide treatment initiation after NG diagnosis. Herein, we developed a NAAT-based phenotypic viability-AST for ciprofloxacin and ceftriaxone against NG (vAST-NG) (Figure 1). By using a membrane-impermeable DNA crosslinker dye, propidium monoazide (PMA, 1), we can differentiate NG strains that are sensitive or susceptible to ciprofloxacin and ceftriaxone from those that are resistant (Scheme 1). Using a logistic regression model, we provide proof of concept that our assay can discriminate between susceptible and resistant strains solely based on the difference between the nucleic acid amplification of the treated and control cells.

**Figure 1.**
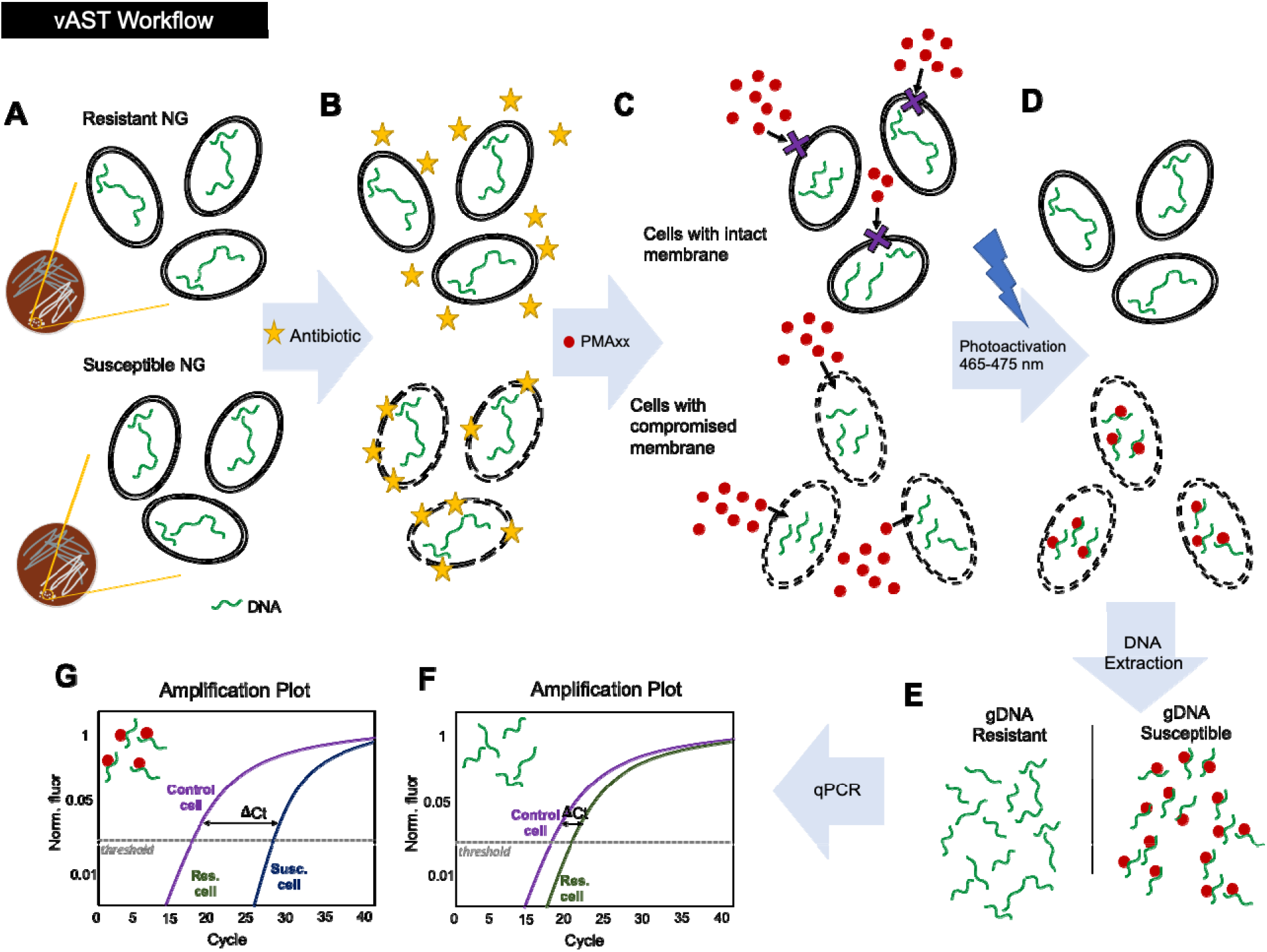
Workflow of vAST-NG for the discrimination of ciprofloxacin and ceftriaxone susceptible cells. (A) Resistant and susceptible WHO reference NG strains were grown overnight on chocolate agar. (B) Cells in GW medium suspensions were treated for 30 min. with antimicrobials (either ciprofloxacin or ceftriaxone) at varying concentrations according to CDC-recommended guideline for both drugs. (C) Cells were then treated with DNA-crosslinker dye, PMAxx™ for 10 min., followed by a 15-min photoactivation using PMALite LED light source (Biotium). (D) After photoactivation, PMAxx™ binds to the DNA of membrane-compromised cells. (E) Genomic DNA of NG cells were extracted using DNA Mini Kit (Qiagen, Australia). (F) Quantitative PCR using TaqMan polymerization reaction was performed for these cells. The nucleic acid amplification plot for resistant strains would typically be very similar to untreated control NG strains, whereas (G) The nucleic acid amplification plot for susceptible strains would show a high cycle threshold value due to the amplification inhibition by PMAxx-bound DNA.

**Scheme 1.**
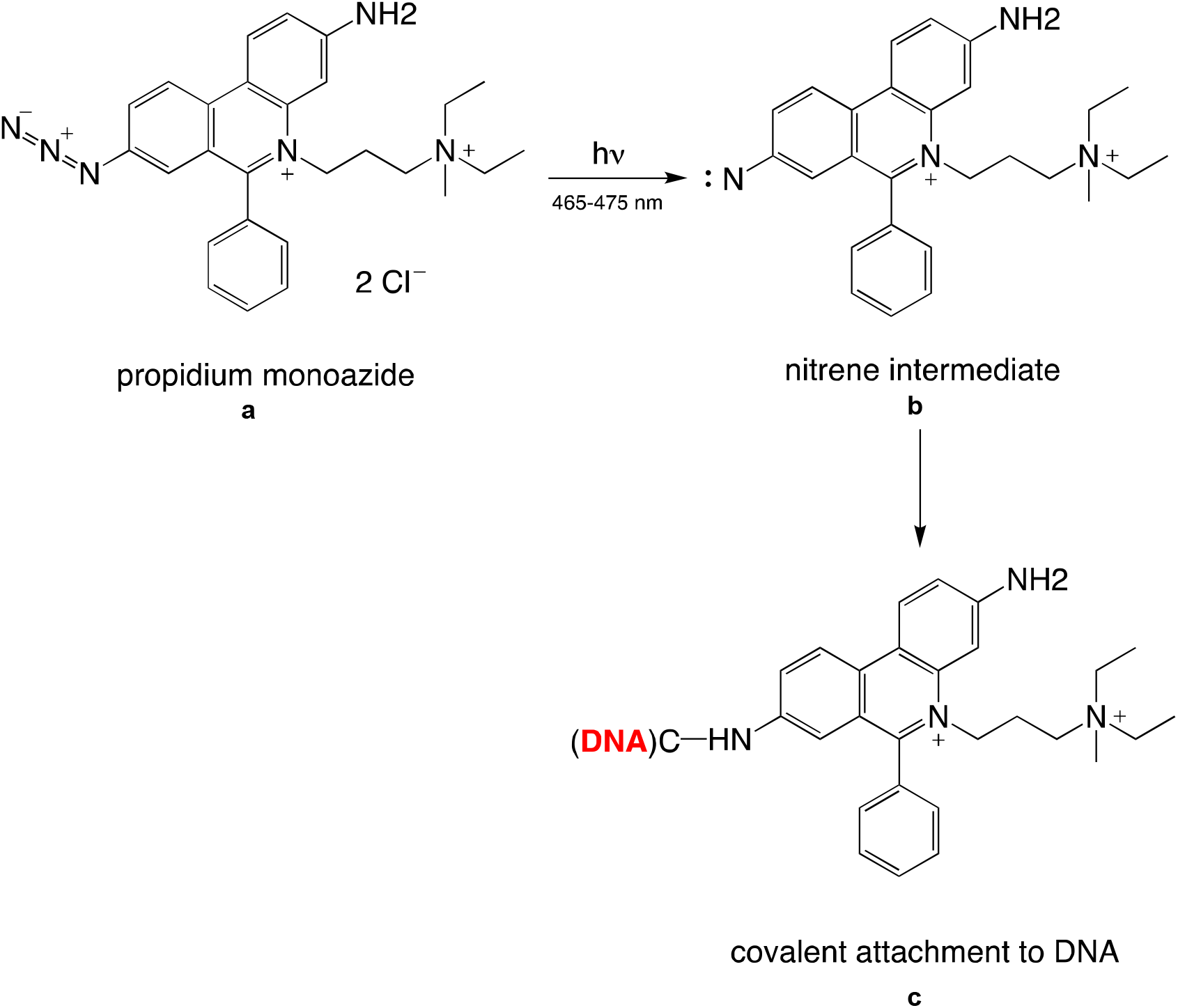
Mechanism of propidium monoazine DNA intercalation^a^. ^*a*^ (a) propidium monoazide, PMA; (b) nitrene intermediate; (c) covalent attachment of PMA to DNA structure modified from van Frankenhuyzen *et al.* (2011) ^16^

## Results

### Viability PCR of ceftriaxone and ciprofloxacin-treated *Neisseria gonorrhoeae* strains

WHO-certified NG reference strains^17^ (Supplemental Table 1) with varied antibiotic susceptibility were treated with ceftriaxone (fourteen strains; 0 to 2 μg/mL) and ciprofloxacin (nine strains; 0 to 8 μg/mL) for 30 min. We confirmed, based on CDC-recommended guideline, using in-house agar dilution and ETEST that the strains used in this study has MICs that concords with the reported MIC (Supplementary Table 2).^17^ After antibiotic incubation, samples were treated with commercially-available DNA crosslinker, PMAxx™ that contains a viability discriminator, propidium monoazide (PMA) dye. Genomic DNA of the NG samples were extracted from the cells and then amplified using quantitative TaqMan qPCR. NG-specific DNA marker for *PorA* pseudogene were used to avoid misclassification.^18^ A set of samples that lacks PMAxx™ were analyzed the same way.

The qPCR cycle threshold number (Ct) was compared between control (untreated) cells and cells treated with either ciprofloxacin or ceftriaxone of varying concentrations. PMAxx™ dye was added to heat-lysed cells as a quality control confirming that the dye can penetrate only to membrane-compromised cells. A significantly higher Ct value was observed for the heat-lysed cells compared to untreated cells in samples treated with DNA-crosslinker dye PMAxx (+). Heat-lysed and untreated samples were indistinguishable when the crosslinker dye is not used, *i.e.,* in PMAxx(-) samples (Supplementary Figure 1). We compared the difference in the Ct value of the ceftriaxone-treated and untreated cells in both resistant and susceptible strains (Figure 2A-B, Supplementary Table 3-4). In general, the PMAxx(-) samples (blue bar) showed no difference in absolute ΔCt across the concentrations while the PMAxx (+) samples (grey bar) showed an increase in absolute ΔCt with increasing ceftriaxone concentration. This dose-dependent effect is more prominent in the susceptible cells (MIC < 0.125, Figure 2B) than in resistant cells (MIC > 0.125, Figure 2A). A higher Ct value was also observed in the PMAxx(+) ceftriaxone-treated susceptible samples at concentration beyond the breakpoint MIC of 0.025 μg/mL in a dose-dependent manner (Figure 2A, grey bars). This increase in Ct value was not observed in PMAxx(-) samples (blue bars).

**Figure 2.**
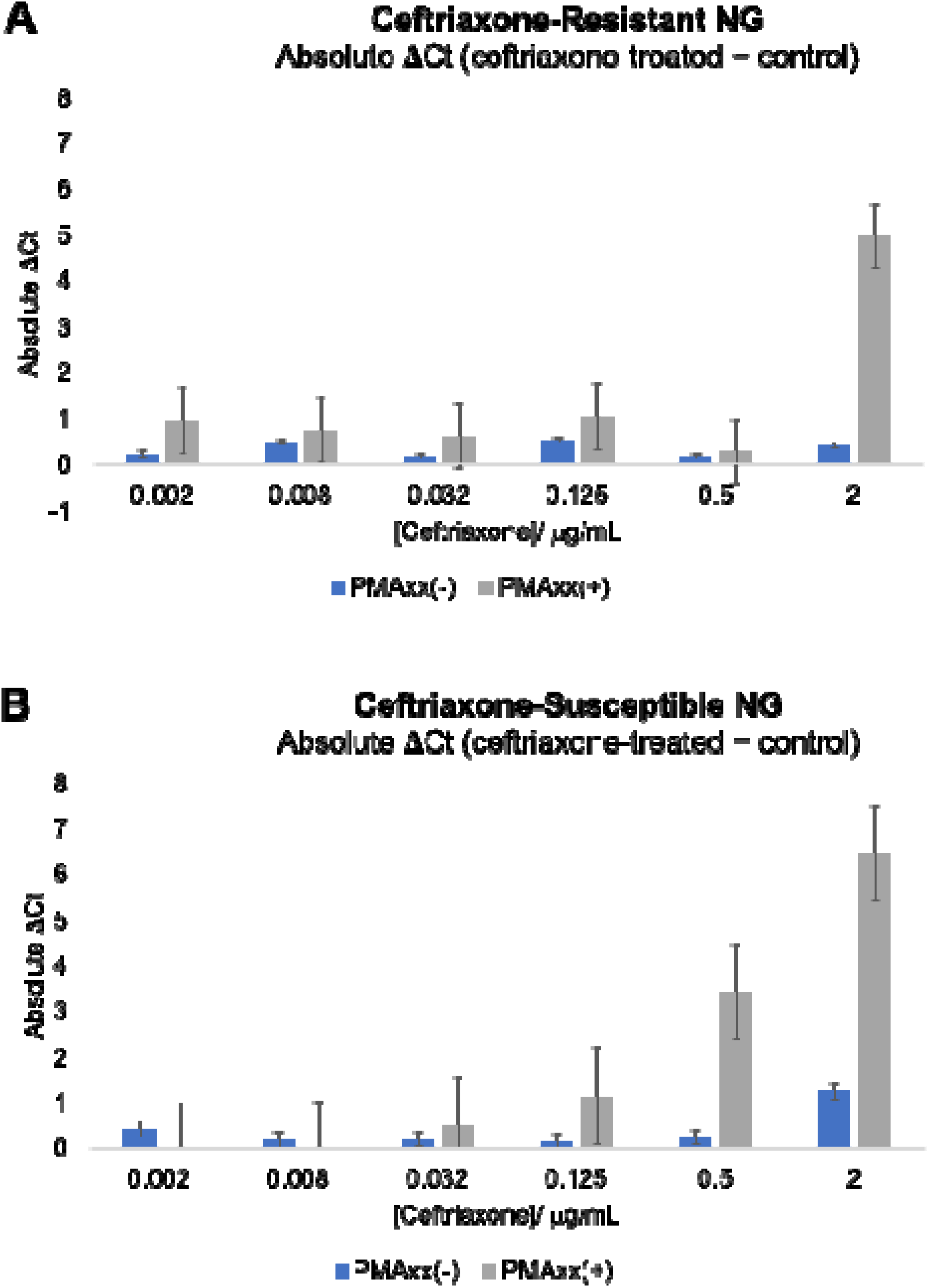
Comparison between the DNA amplification cycle thresholds of PMAxx-treated and untreated genomic DNA samples from 14 WHO reference NG strains. (A) The absolute difference between the Ct value of PMAxx-treated or untreated control live cells (not treated with ceftriaxone) and ceftriaxone-treated cells of all ceftriaxone-resistant strains (WHO X, L, Y, Z, MIC > 0.125 μg/mL). (B) The absolute difference between the Ct value of PMAxx-treated or untreated control live cells and ceftriaxone-treated cells of all ceftriaxone-susceptible strains (WHO F, O, G, K, M, P, W, V, N, U, MIC < 0.125 μg/mL). Error bars represent standard deviation of the mean.

### Assay digital PCR

The vAST-NG assay was performed on a digital PCR instrument to evaluate the assay’s feasibility on this platform. The same protocol was tested for two NG strains on a QuantStudio Absolute Q digital PCR platform versus qPCR (Supplementary Table 5). Our preliminary result demonstrated that when using target DNA copy number as readout from digital PCR, a higher magnitude of difference was observed between the treated versus untreated sample as compared to Ct value difference from qPCR. This result demonstrates the potential translation of the assay to a digital PCR platform.

### Determining *N. gonorrhoeae* susceptibility to ceftriaxone and ciprofloxacin

A logistic regression model trained on the natural log transformed mean absolute difference in Ct values determined after 30 minutes of exposure to the antibiotic performed with an AUC of 0.93 (Figure 3A). The natural log transformed mean absolute difference in Ct under concentrations 0.002 and 0.5 μg/mL were significant predictors of susceptibility with odd ratios (OR) of 47.78 [95% confidence interval (CI): 2.62 – 73434.22] for 0.002 μg/mL and 11.55 [95% CI: 1.42 – 796.13] for 0.5 μg/mL. The ceftriaxone and ciprofloxacin combined model had a categorical agreement for susceptibility was 84.6% with a major error of 15.4% (Figure 3B). Similarly, the categorical agreement for resistance to the antibiotic was 80% with a very major error of 20%.

**Figure 3.**
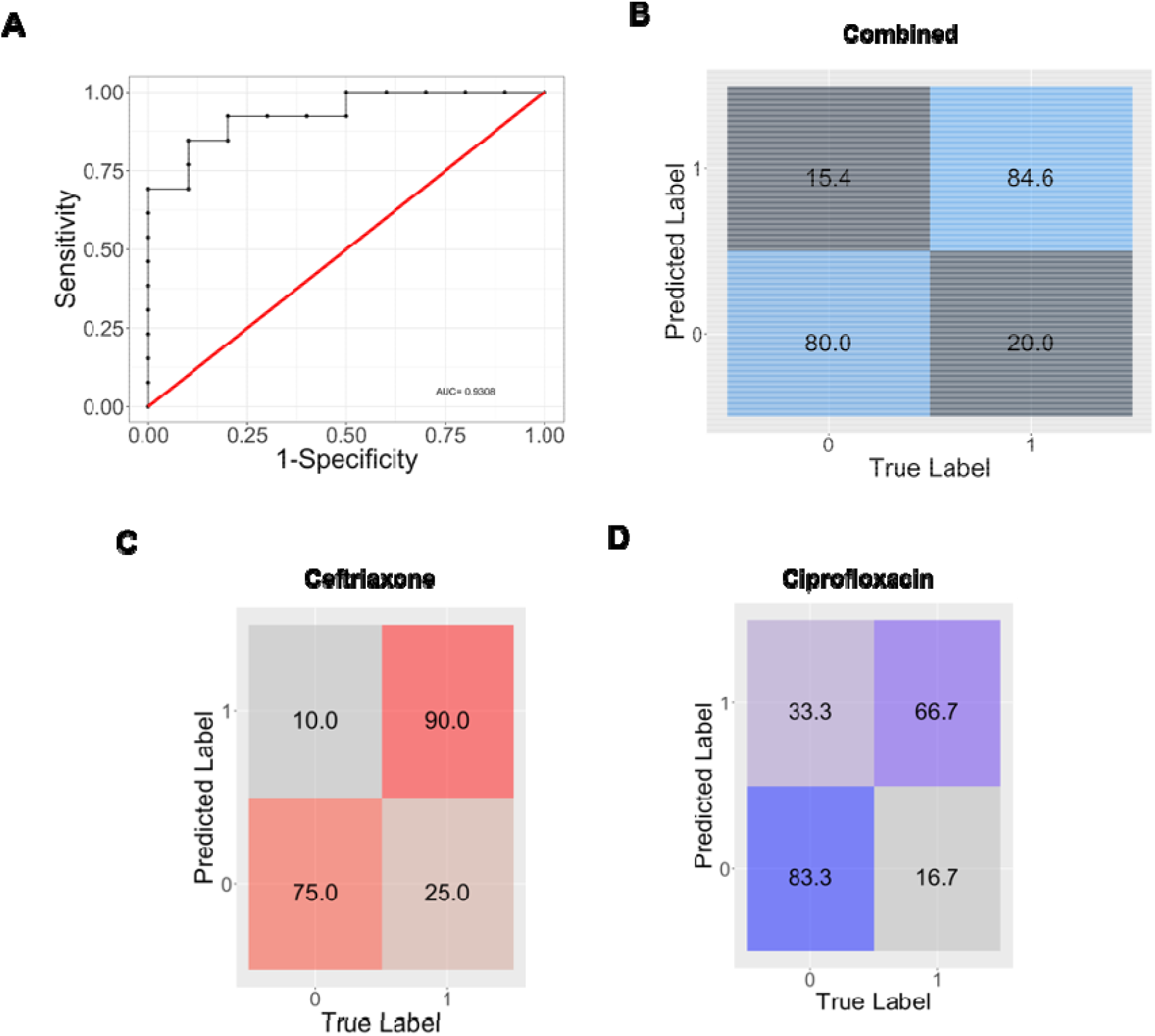
Performance of the combined logistic regression model. A. Receiver operator curve of the ceftriaxone and ciprofloxacin combined logistic regression model on the mean of the log2 transformed absolute Ct difference between live (control) and treated cells had an AUC of 0.93. (B) Confusion matrix of the combined model with >80% accuracy in classifying susceptibility to the two antimicrobials. (C) Confusion matrix for ceftriaxone-treated NG samples. (D) Confusion matrix for ciprofloxacin-treated NG samples.

When surveying the performance against each antibiotic separately, the model predicted 9/10 and 3/4 susceptible and resistant strains respectively to ceftriaxone (Figure 3C) and 2/3 and 5/6 susceptible and resistant strains respectively to ciprofloxacin (Figure 3D). We further validated the model against the individual replicates. For ceftriaxone, the model accurately predicted 23/27 susceptible and 9/12 resistant replicates with 100% accuracy in prediction for 6/10 susceptible strains. For ciprofloxacin, the model accurately predicted 6/9 susceptible and 10/18 resistant replicates. Hence, we confirmed that the susceptibility information generated using our protocol concords with the reported standard ^17^.

We then assayed the ability of the model to predict susceptibility agnostic of the inoculum size. We tested three replicates of two strains (WHO O susceptible and WHO X resistant strains to ceftriaxone) with an inoculum size of 10^2^ CFU/mL and repeated the viability PCR to collect Ct values after 30 minutes of exposure to different concentrations of antibiotic. While the combined ceftriaxone and ciprofloxacin model misclassified all replicates as susceptible, a logistic regression model trained solely on ceftriaxone data for the 10^5^ CFU/mL inoculum size was able to successfully classify 3/3 susceptible replicates and 2/3 resistant replicates for the lower inoculum size (Figure 4). Accurate classification of susceptibility was achieved agnostic of the inoculum size and once again, the difference in Ct values obtained at the concentrations of 0.002 and 0.5 μg/mL influenced the model outcome.

**Figure 4.**
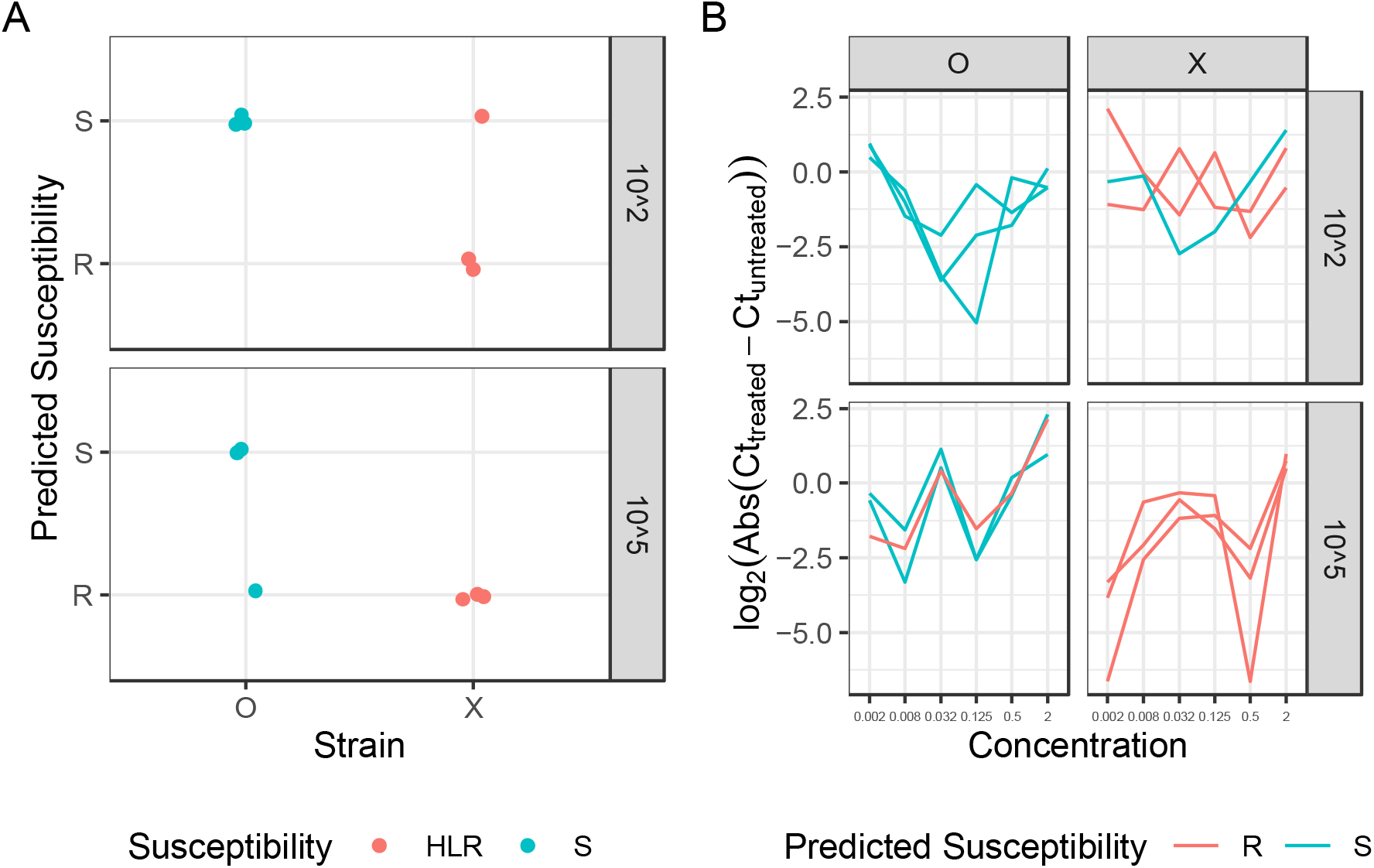
Accurate classification of susceptibility agnostic of inoculum size. A. Logistic regression model trained on ceftriaxone treated NG with a 10^5^ CFU/mL inoculum size successfully classified majority of the replicates (2/3) accurately thereby classifying strain O as susceptible and strain X resistant when assayed at an inoculum size of 10^2^ CFU/mL. B. Processed data for the 3 replicates of the two strains used to test the algorithm. Values obtained at concentrations of 0.002 and 0.5 μg/mL influenced the susceptibility prediction by the model.

## DISCUSSION

Herein, we described a rapid phenotypic AST based on MIC against 2 NG-relevant antimicrobials –ceftriaxone and ciprofloxacin. We leveraged the sensitivity, specificity and speed of mainstay qPCR methods routinely performed for NG detection in the clinic and develop an AST test that can potentially be integrated into mainstream NG PCR assays. We demonstrated that with only 30-minutes antimicrobial incubation on 14 WHO reference NG strains, we could achieve dose-response output based on the qPCR cycle threshold (Ct). Using a logistic regression model, an AUC of 0.93 with an 84.6% categorical agreement for susceptibility was obtained for both antimicrobials.

Due to the lack of new antimicrobials in the pipeline, repurposing existing antimicrobials is the only option for treating ceftriaxone-resistant cases. The GISP 2019 reports around 35.4% of NG cases in the US are still susceptible to ciprofloxacin (0.1% of NG isolates are resistant to ceftriaxone) ^5^. While ceftriaxone treatment failure due to resistance-related reasons has not been identified in the United States, it is beginning to appear in Asia, Australia, and the United Kingdom ^9,14,19^ Emerging cases of NG isolates with alert values (MIC ≥ 0.125 μg/mL) justify the need for increased screening measures to control the potential importation of resistance mutation and its rapid transmission within a community. While rapid ciprofloxacin resistance tests NG based on genotypic markers have been developed, rapid phenotypic tests are currently unavailable ^20,21^.

Our proposed use of vAST-NG can be found in clinical workflow outlined in Figure 5. Two urogenital swabs will be collected from each NG suspected patient. Once NG diagnosis is made using swab 1, a second swab will be used. To inform susceptibility, we will split swab 2 in culture media into 3 aliquots and challenge each with antimicrobials (ciprofloxacin and ceftriaxone) at CLSI breakpoint concentrations (see Methods), followed by treatment with a photoreactive DNA-crosslinker dye containing propidium monoazide (marketed as PMAxx™). Propidium monoazide (PMA) has previously been used to assess the viability of various microorganisms in different contexts ^22–25^. Its use in antibiotic susceptibility tests with qPCR has been demonstrated in *Mycobacterium tuberculosis,* but not in NG ^22^. Aside from the dye, we also used an NG-specific PCR primers and probes to minimize the chance of cross-reactions ^15^. The assay readout is in the form of a categorical susceptibility (susceptible or resistant) indicated by the number of amplification cycles required to detect NG DNA (cycle threshold, or ‘Ct’). A higher Ct value implies a higher cell membrane permeability and hence a higher level of DNA intercalation by PMA dye that leads to amplification inhibition. Using this principle, an antimicrobial dose-response effect can be assessed using vAST-NG after a 30 min antimicrobial incubation.

**Figure 5.**
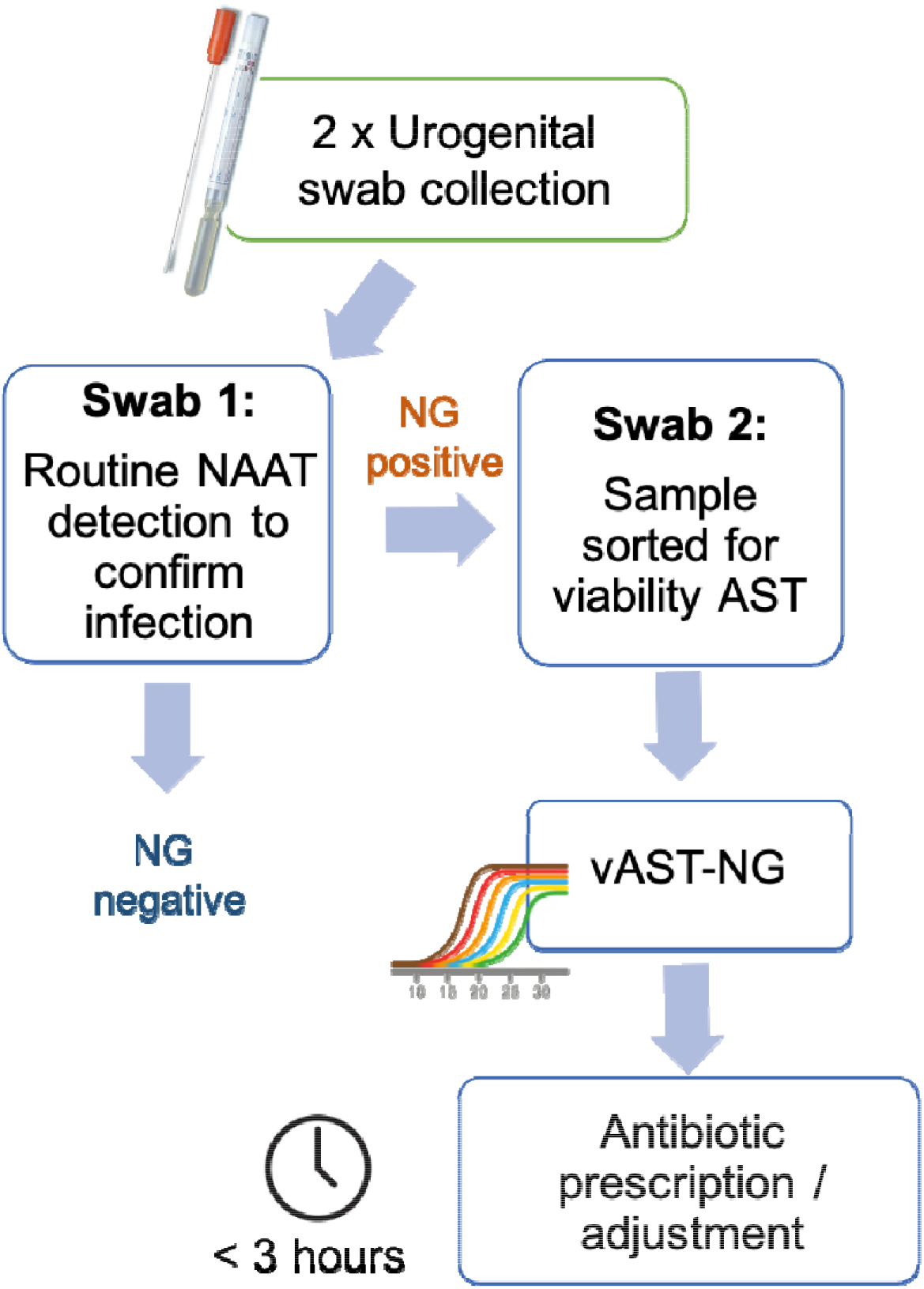
Proposed clinical workflow for vAST-NG. Two urogenital swabs will be collected from patients with suspected NG infections. The first swab sample will be used for routing NAAT detection. If positive, the second swab will be used for vAST-NG assay. The result of the assay, in the form of susceptibility information for both ciprofloxacin and ceftriaxone, will guide antibiotic prescription or adjustment.

In addition to a qPCR, we have also tested our assay on a digital PCR platform using the same sample preparation procedure without significant modifications (Supplementary Table 5). Translating the assay to a digital platform could potentially improve assay sensitivity, particularly in samples with low NG load.

Determining antimicrobial susceptibility for NG isolates is important for both surveillance and clinical management. In the era where NG diagnosis is predominately made *via* nucleic-acid-based molecular testing (e.g. PCR), standard phenotypic antimicrobial susceptibility testing (AST), which is time-consuming (24-48 hrs.), laborious, and requires viable cultures, is not routinely performed. Methods that rely on detecting a single mutation as a marker of resistance have been explored, however ongoing monitoring of resistance patterns is required^15,20^. Relying on a single resistance marker can easily blindside complex and evolving resistance mechanisms ^13^. The use of high-throughput genomic analysis, such as whole genome sequencing (WGS), has been suggested as a suitable alternative to address this limitation. However, WGS could only be used to detect known resistance determinants and cannot reliably provide minimum inhibitory concentration necessary for surveillance purposes. Misclassification of susceptibility by relying on the absence of known resistance markers may lead to very major errors (VME). Therefore, tests that ultimately rely on phenotypic cues could give greater confidence in capturing true susceptibility regardless of genotypic changes.

Phenotypic AST for NG directly from clinical samples without culture expansion, on the other hand, has its own challenges. International guidelines for reference phenotypic ASTs, such as broth microdilution, standardize inoculum size to 2 to 8 x 10^5^ CFU/mL. This pathogen load is typical for NG samples collected from the urine, vaginal swab, and anorectal swabs of both asymptomatic and symptomatic men and women.^26^ The NG pathogen load for an oropharyngeal sample tends to be at a lower, 10^3^ CFU/mL range.^26^ Deviation from the standard guideline with variable inoculum size input makes analysis challenging and could risk potential inoculum effect.^27,28^ On the other hand, this requirement can be problematic for slow-growing organisms such as NG where the need for culture can significantly increase assay turnaround. We circumvented this problem by utilizing a machine learning algorithm based on logistic regression to classify the resistant/susceptible probability based on our test samples. To demonstrate this versatility, we tested our assay using one ceftriaxone-resistant (WHO X) and one ceftriaxone-susceptible (WHO O) NG at 10^2^ CFU/mL done in triplicates. We found that using the logistic regression model trained on Ct values from 10^5^ CFU/mL inoculum size, we were able to successfully classify all 3 replicates of the susceptible strains and 2 out of 3 replicates of the resistant strains at low inoculum size of 10^2^ CFU/mL. Measurements using a variety of strains and cell concentrations will be needed to account for a more robust training model.

## CONCLUSION

The vAST-NG assay is designed to deliver clinically relevant phenotypic antimicrobial susceptibility information for NG. The assay serves two purposes: (1) to enable an evidence-based repurposing of antimicrobials for NG treatment, and (2) to monitor the population of NG strains that are resistant to ceftriaxone and anticipate any potential outbreak of ceftriaxone-resistant NG strains. The assay provides susceptibility information by measuring cell viability. Since this method relies on quantifiable Ct values between samples, the readouts are both phenotypic and objective.

Future development of this assay will involve testing on a larger sample size to improve assay accuracy. We envision vAST-NG to be easily incorporated into any commercially available PCR assay (both quantitative and digital) for NG diagnosis that combines both species identification (ID) with phenotypic antimicrobial susceptibility testing (AST). This assay could potentially serve as a reflex test immediately after a positive NG diagnosis by routine POC NAAT. Since NAAT such as qPCR is already a mainstay in many clinical laboratories for NG diagnosis, further development could allow immediate translation without specialized laboratory training. With the availability of adjacent automated sample preparation instrumentation, future development of this approach can be performed with minimal hands-on time and requires fewer consumables.

Our current report serves as a proof-of-concept for the use of viability PCR as a method to rapidly inform phenotypic AST in NG. The results outlined in this report are based on a limited dataset acquired from 14 WHO reference NG strains using a uniformed inoculum size. Future effort will focus on two areas. Firstly, increasing sample size to train the vAST-NG logistic regression model and improve assay prediction. Secondly, establishing the assay limit of detection to reach a clinically relevant cell density in both symptomatic and asymptomatic gonorrhea cases.

## METHODS

### Microorganisms and culturing

Fourteen WHO (F, G, K, L, M, N, O, P, U, V, W, X, Y, Z) reference NG strains obtained from Johns Hopkins Medicine, Department of Pathology, Division of Medical Microbiology were used for the study (Supplementary Table 1). These Isolates were cultured from −80 °C glycerol stocks on Chocolate II Agar (BD BBL™ Prepared Plated Media: GC II Agar with Hemoglobin and IsoVitaleX™, #B21267X, Becton Dickinson, United States). Plates were incubated at 35 °C, in 5% CO_2_ for 24 h.

### Cell preparation and antibiotic treatment

For each NG reference strains, overnight colonies were re-suspended in GW broth to an optical density equal to that of a 0.5 MacFarland standard or equivalent to 10^8^ colony forming unit (CFU) per mL. The cell suspension was diluted at 1:1000 ratio in GW broth. Eight 2 mL of 10^5^ CFU/mL NG suspension in culture tubes were prepared. Six of the eight tubes were treated with 0.002, 0.008, 0.032, 0.125, 0.5 and 2 μg/mL of ceftriaxone. Two of the remaining 2 mL NG suspensions serve as positive ‘Live’ and negative ‘Dead’ controls. All eight tubes were incubated at 35 °C, in 5% CO_2_ for 2 h with constant shaking (200 rpm). After 30 min., the 8 x 2 mL bacteria suspension were and transferred into 1.5 mL microcentrifuge tubes at equal volume (16 x 1 mL). Eight of the 1 mL bacteria suspension were used for treatment with viability dye, while the other eight serve as controls. Negative Dead control cells (2 x 1 mL) were incubated at 90 °C for 5 min to lyse cells. Stock Ciprofloxacin solution (8 mg/mL) was prepared by dissolving ciprofloxacin (8 mg) in HCl (20 μL) and GW broth to a total volume of 1 mL.

### Viability discrimination assay

PMAxx™ DNA-intercalating dye (Biotium, United States) was used to distinguish between viable and nonviable cells. Manufacturer protocol was followed with slight modification. PMAxx™ DNA-intercalating dye (1.25 mL of 20 mM stock) was added to eight of the 1 mL bacteria suspension, followed by a 10 min incubation in dark. After the incubation, a 15 min photoactivation using the PMA-Lite™ LED Photolysis Device was performed as recommended by the PMAxx™ manufacturer (Biotium, Inc., United States). After 15 min, the cells were centrifuged at 8000 rpm for 10 min. Supernatant was carefully discarded, followed by cell lysis and genomic DNA extraction.

### Genomic DNA extraction

Genomic DNA extraction was carried out on a QIACube Connect using a QIAamp® DNA Mini Kit (Qiagen). A 50 μL elution volume was used at the final stage of the extraction. In short, cell lysis was performed at 56 °C for 30 min followed by a 10 min incubation at 70 °C for 10 min using the recommended Qiagen buffers. At the end of the extraction, 150 μL of genomic DNA from 16 NG samples were obtained.

### Real-Time quantitative PCR with Taqman probe

Real-time quantitative PCR amplification was performed using the TaqMan™ Fast Advanced Master Mix (Applied Biosystems, United States), previously-validated^18^ TaqMan dual-labeled probe (FAM-CGCCTATACGCCTGCTACTTTCACGC-BHQ^1^) and primers (papF: CAGCATTCAATTTGTTCCGAGTC, papR:GAACTGGTTTCATCTGATTACTTTCCA) targeting the *PorA* genes that were ordered from the Stanford PAN Facility. The reaction was performed in a final volume of 21 μL containing of 2 μL of each primer and probe (10 μM), 10 μL of TaqMan Fast Advanced Master Mix, and 5 μL of genomic DNA extract. The amplification reaction was conducted using a Rotor-Gene-Q PCR Cycler Real-Time PCR system (Qiagen, Australia). The cycling parameters were as follows: initial denaturation step at 95°C for 20 s, followed by 40 cycles of denaturation at 95°C for 1 s and annealing / extension at 60°C for 20 s.

### Data analysis

The expression of each candidate marker was automatically detected in the green channel (FAM) by the Rotor-Gene Q Series software. A threshold of 0.02 was set to obtain accurate cycle threshold (Ct) values for each isolate. All amplification reactions were run in triplicates. To calculate the ΔCt value, control isolates that are not treated with PMAxx™ dye were used to normalize the potential difference between untreated control and antibiotic-treated samples. Fold change in gene expression was calculated as FC = 2^-ΔΔCt^. The absolute difference between the Ct value for living and treated cells was log2 transformed and the mean of three replicates was used to develop a logistic regression model using glm in R.

### Antimicrobial susceptibility testing

The CDC reported MICs were confirmed in-house by performing agar dilution, the gold standard antimicrobial susceptibility test for NG recommended by the CDC. https://www.cdc.gov/std/gonorrhea/lab/agar.htm. Viability qPCR results obtained from the assay was validated within MIC ± 1 log2 dilution of the reported MIC against this standard.

## Acknowledgements

We also thank Justin Hardick for the experimental advice and the Johns Hopkins Medicine, Department of Pathology, Division of Medical Microbiology for the clinical isolates.

