## Supplementary Material for "Rapid molecular phenotypic antimicrobial susceptibility test for *Neisseria gonorrhoeae* based on propidium monoazide viability PCR"

**Author Affiliations:**

**Supplementary Table 1**. WHO *N. gonorrhoeae* reference strains and their reported MIC used in this study[24]

| **No** | **Strains** | **Ceftriaxone** | **Ciprofloxacin** |
| --- | --- | --- | --- |
| 1 | WHO F | <0.002 (S) | S (0.004) |
| 2 | WHO G | 0.008 (S) | LLR (0.125) |
| 3 | WHO K | 0.064 (S) | HLR (>32) |
| 4 | WHO L | 0.25 (LLR) | HLR (>32) |
| 5 | WHO M | 0.016 (S) | R (2) |
| 6 | WHO N | 0.004 (S) | R (4) |
| 7 | WHO O | 0.032 (S) | 0.008 (S) |
| 8 | WHO P | 0.004 (S) | S (0.004) |
| 9 | WHO U | 0.002 (S) | S (0.004) |
| 10 | WHO V | 0.064 (S) | HLR (>32) |
| 11 | WHO W | 0.064 (S) | HLR (>32) |
| 12 | WHO X | 2 (HLR) | >32 (HLR) |
| 13 | WHO Y | 1 (HLR) | HLR (>32) |
| 14 | WHO Z | 0.5 (LLR) | HLR (>32) |

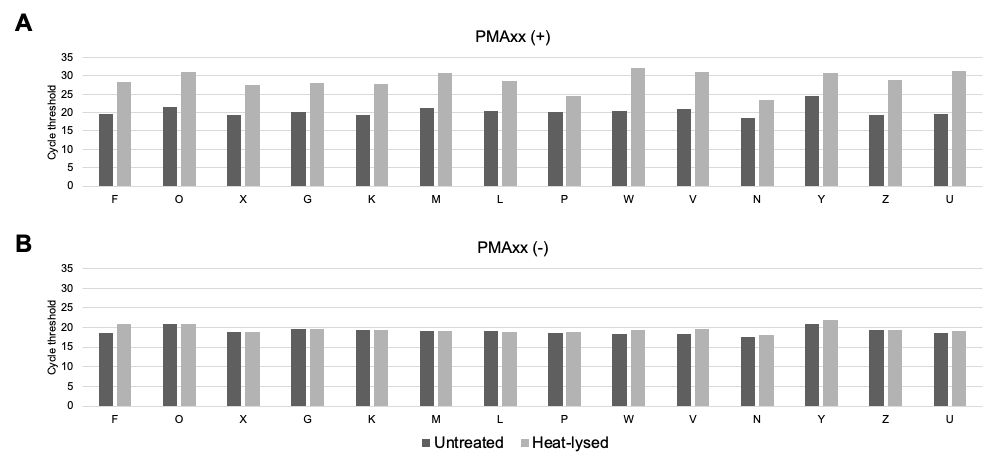

**Supplementary Figure 1.** Cycle threshold comparison between heat-lysed and untreated NG WHO reference strains**.** (A) PMAxx DNA-crosslinker dye was added to the sample. (B) PMAxx DNA-crosslinker dye was not used.

**Supplementary Table 2.** Validation of ceftriaxone MIC for WHO *N. gonorrhoeae* reference strains using in-house agar dilution and ETEST based on CDC guideline (www.cdc.gov)

| **No** | **Strains** | **Reported S/I/R** | **Experimental  Agar Dilution** | **Experimental ETEST** | **Susc. category using  viability PCR** | **Agreement  with CLSI Category** |
| --- | --- | --- | --- | --- | --- | --- |
| 1 | WHO F | S | 0.008 | <0.002 | S | Y |
| 2 | WHO G | S | 0.016 | 0.008 | S | Y |
| 3 | WHO K | S | 0.064 | 0.064 | S | Y |
| 4 | WHO L | LLR | 0.25 | 0.125 | R | Y |
| 5 | WHO M | S | 0.032 | 0.004 | S | Y |
| 6 | WHO N | S | 0.008 | 0.006 | S | Y |
| 7 | WHO O | S | 0.032 | 0.008 | S | Y |
| 8 | WHO P | S | 0.032 | 0.006 | S | Y |
| 9 | WHO U | S | 0.008 | <0.002 | S | Y |
| 10 | WHO V | S | 0.032 | 0.016 | S | Y |
| 11 | WHO W | S | 0.064 | 0.032 | S | Y |
| 12 | WHO X | HLR | 1 to 2 | 1 | R | Y |
| 13 | WHO Y | HLR | 0.125 | 1.5 | R | Y |
| 14 | WHO Z | LLR | 0.25 | 0.5 | R | Y |
| 15 | F18 | S | 0.032 | 0.016 | S | Y |
| 16 | SPJ15 | S | 0.016 | 0.012 | S | Y |
| 17 | SPL4 | LLR | 0.25 | 0.19 | R | Y |

**Supplementary Table 3.** Absolute Ct difference between ceftriaxone treated and untreated resistant NG with and without Pmaxx.

| WHO Strains | Concentration | w/Pmaxx | w/o Pmaxx |
| --- | --- | --- | --- |
|  | 0.002 | 0.05 | 1.53 |
| L | 0.008 | 0.07 | 1.78 |
|  | 0.032 | 0.94 | 1.56 |
|  | 0.125 | 0.15 | 0.94 |
|  | 0.5 | 2.03 | 0.06 |
|  | 2 | 5.85 | 0.23 |
|  | 0.002 | 0.55 | 0.09 |
|  | 0.008 | 0.29 | 0.45 |
| X | 0.032 | 0.35 | 0.06 |
|  | 0.125 | 0.33 | 0.3 |
|  | 0.5 | 0.83 | 0.24 |
|  | 2 | 5.81 | 0.77 |
|  | 0.002 | 0.19 | 0.09 |
| Y | 0.008 | 1.62 | 0.45 |
|  | 0.032 | 0.85 | 0.18 |
|  | 0.125 | 1.14 | 2.97 |
|  | 0.5 | 0.9 | 1.14 |
|  | 2 | 3.01 | 1.24 |
|  | 0.002 | 0.61 | 0.44 |
| Z | 0.008 | 0.71 | 0.65 |
|  | 0.032 | 0.34 | 0.08 |
|  | 0.125 | 0.04 | 0.75 |
|  | 0.5 | 0.73 | 0.71 |
|  | 2 | 6.25 | 0.15 |

**Supplementary Table 4.** Absolute Ct difference between ceftriaxone treated and untreated susceptible NG with and without Pmaxx.

| WHO Strains | Concentration | w/Pmaxx | w/o Pmaxx |
| --- | --- | --- | --- |
|  | 0.002 | 0.9 | 1.31 |
|  | 0.008 | 0.39 | 1.51 |
|  | 0.032 | 1.72 | 1.48 |
| F | 0.125 | 3.97 | 0.54 |
|  | 0.5 | 7.18 | 1.88 |
|  | 2 | 6.85 | 1.27 |
|  | 0.002 | 0.21 | 0.52 |
|  | 0.008 | 0.02 | 0.3 |
| K | 0.032 | 0.94 | 0.34 |
|  | 0.125 | 1.49 | 0.05 |
|  | 0.5 | 2.78 | 2.03 |
|  | 2 | 7.25 | 1.24 |
|  | 0.002 | 0.12 | 1.49 |
| G | 0.008 | 0.45 | 2.22 |
|  | 0.032 | 0.58 | 1.5 |
|  | 0.125 | 0.77 | 1.57 |
|  | 0.5 | 6.55 | 0.68 |
|  | 2 | 5.43 | 0.93 |
|  | 0.002 | 0.63 | 0.41 |
| O | 0.008 | 0.09 | 0.65 |
|  | 0.032 | 0.37 | 0.65 |
|  | 0.125 | 1.15 | 0.59 |
|  | 0.5 | 3.03 | 0.52 |
|  | 2 | 6.45 | 1.29 |
|  | 0.002 | 0.31 | 0.15 |
| W | 0.008 | 0.78 | 0.17 |
|  | 0.032 | 0.83 | 0.44 |
|  | 0.125 | 0.46 | 0.12 |
|  | 0.5 | 1.96 | 0.56 |
|  | 2 | 5.92 | 0.28 |
|  | 0.002 | 1.55 | 0.97 |
| M | 0.008 | 0.85 | 0.78 |
|  | 0.032 | 0.51 | 0.41 |
|  | 0.125 | 0.11 | 0.82 |
|  | 0.5 | 1.19 | 0.71 |
|  | 2 | 6.21 | 0.11 |
|  | 0.002 | 0.17 | 0.58 |
|  | 0.008 | 0.15 | 0.49 |
| P | 0.032 | 1.3 | 0.01 |
|  | 0.125 | 2.61 | 0.04 |
|  | 0.5 | 4.74 | 0.56 |
|  | 2 | 6 | 1.32 |
|  | 0.002 | 0.45 | 0.19 |
| V | 0.008 | 1.39 | 0.36 |
|  | 0.032 | 1.49 | 0.33 |
|  | 0.125 | 1.41 | 0.36 |
|  | 0.5 | 1.3 | 0.4 |
|  | 2 | 4.65 | 1.73 |
|  | 0.002 | 0.03 | 0.77 |
|  | 0.008 | 0.37 | 0.35 |
| N | 0.032 | 0.43 | 0.04 |
|  | 0.125 | 1.94 | 0.17 |
|  | 0.5 | 3.13 | 0.57 |
|  | 2 | 5.98 | 1.23 |
|  | 0.002 | 0.36 | 0.14 |
|  | 0.008 | 0.07 | 0.08 |
| U | 0.032 | 0.97 | 0 |
|  | 0.125 | 2.17 | 0.54 |
|  | 0.5 | 7.52 | 1.53 |
|  | 2 | 8.8 | 1.38 |

**Supplementary Table 5.** Comparison of vAST-NG amplification result between dPCR and qPCR result

| Strains | WHO O (Susceptible | | WHO X (Resistant) | |
| --- | --- | --- | --- | --- |
|  | qPCR (Ct) | dPCR (copies/uL) | qPCR (Ct) | dPCR (copies/uL) |
| Live (untreated) | 27.37 | 333.99 | 27.35 | 503.316 |
| 0.002 mg/L | 27.92 | 311.094 | 27.72 | 395.352 |
| 2 mg/L | 31.71 | 27.558 | 29.23 | 160.758 |
